# Persistent Disruptions in Prefrontal Connectivity Despite Behavioral Rescue by Environmental Enrichment in a Mouse Model of Rett Syndrome

**DOI:** 10.1101/2025.02.10.637474

**Authors:** Sofie Ährlund-Richter, Jonathan Harpe, Giselle Fernandes, Ruby Lam, Mriganka Sur

## Abstract

Rett Syndrome, a neurodevelopmental disorder caused by loss-of-function mutations in the *MECP2* gene, is characterized by severe motor, cognitive and emotional impairments. Some of the deficits may result from changes in cortical connections, especially downstream projections of the prefrontal cortex, which may also be targets of restoration following rearing conditions such as environmental enrichment that alleviate specific symptoms. Here, using a heterozygous *Mecp2^+/−^* female mouse model closely analogous to human Rett Syndrome, we investigated the impact of early environmental enrichment on behavioral deficits and prefrontal cortex connectivity. Behavioral analyses revealed that enriched housing rescued fine motor deficits and reduced anxiety, with enrichment-housed *Mecp2^+/−^* mice performing comparably to wild-type (WT) controls in rotarod and open field assays. Anatomical mapping of top-down anterior cingulate cortex (ACA) projections demonstrated altered prefrontal cortex connectivity in *Mecp2^+/−^* mice, with increased axonal density in the somatosensory cortex and decreased density in the motor cortex compared to WT controls. ACA axons revealed shifts in hemispheric distribution, particularly in the medial network regions, with *Mecp2^+/−^*mice exhibiting reduced ipsilateral dominance. These changes were unaffected by enriched housing, suggesting that structural abnormalities in prefrontal cortex connectivity persist despite behavioral improvements. Enriched housing rescued brain-derived neurotrophic factor (BDNF) levels in the hippocampus but failed to restore BDNF levels in the prefrontal cortex, consistent with the persistent deficits observed in prefrontal axonal projections. These findings highlight the focal nature of changes induced by reduction of MeCP2 and by exposure to environmental enrichment, and suggest that environmental enrichment starting in adolescence can alleviate behavioral deficits without reversing abnormalities in large-scale cortical connectivity.

## INTRODUCTION

Rett Syndrome (RTT) is a severe neurodevelopmental disorder primarily affecting girls and caused by loss-of-function mutations in the X-linked gene which encodes the methyl-CpG-binding protein 2 (MeCP2)(Amir et al., 1999; Chahrour and Zoghbi, 2007). Patients diagnosed with RTT exhibit a range of symptoms including intellectual disabilities, motor dysfunction, autistic features, and abnormalities in their heart rate and breathing(Amir et al., 1999; Buchanan et al., 2019). In addition, RTT patients commonly suffer from devastating general and/or social anxiety, which may present as hyperventilation, panic attacks, inconsolable crying, trembling in the absence of fearful situations, screaming episodes, gaze avoidance and withdrawal from social contact (Buchanan et al., 2019). Neural circuit dysfunction is thought to be a hallmark of RTT with several studies reporting changes in dendritic structure and synaptic function accompanying loss of MeCP2 in mice (Chao et al., 2007; Belichenko et al., 2008; Tropea et al., 2009; Banerjee et al., 2019; Sandweiss et al., 2020). However, the effects of MeCP2 loss on long-range axonal connectivity and its contribution to circuit defects and behavioral abnormalities in RTT are unknown.

Neuroimaging studies in RTT patients show marked reduction in gross cerebral volume particularly in the frontal, parietal and temporal lobes (Carter et al., 2008; Takeguchi et al., 2022), as well as a decrease in the size of white matter tracts (Reiss et al., 1993; Naidu et al., 2001). These changes in the structure and density of neurons and axons suggest that loss of MeCP2 function severely disrupts the structure of neural circuits in the brain. Although MeCP2 is expressed brain-wide, it has become evident that physiological changes observed due to its loss are region-specific and can vary widely between brain regions (Ip et al., 2018). For example, Scholl analyses in postmortem samples of RTT patients show atrophy of dendritic trees in the frontal and motor cortex (Armstrong et al., 1995). Similarly, tracing studies show that axonal fibers in the hippocampus and motor cortex are disorganized in both RTT patients (Belichenko et al., 1994) and Mecp2 null mice (Belichenko et al., 2009). However, no atrophy is observed in the visual cortex of RTT patients (Armstrong et al., 1995). These regional differences in structural alterations due to loss of MeCP2 may underlie alterations in behavioral function whereby motor function is severely reduced but vision appears normal (Armstrong et al., 1995). Further, studies exploring the molecular mechanisms of MeCP2 function have revealed that MeCP2 acts as a transcriptional regulator, altering the focal expression of neurotrophins such as brain-derived neurotrophic factor (BDNF) and insulin-like growth factor-1 (IGF-1), both of which have been implicated in the development of neural circuits (Chen et al., 2003; Chang et al., 2006; Castro et al., 2014; Ip et al., 2018). This highlights the necessity of exploring neural circuit changes across brain regions, especially in areas associated with the behavioral symptoms of RTT. One such region is the prefrontal cortex.

The prefrontal cortex is considered a major cognitive center in the brain, continuously integrating and modulating neuronal representations of internal and external information (Fuster, 2015). This requires the prefrontal cortex to have access to the current behavioral state, and process sensory information in relation to our current needs. Consequently, the prefrontal cortex is highly implicated in the regulation of anxiety, and has been suggested to modify the experience of anxiety to shape behavior and emotion in response to a stressor (Kenwood et al., 2022). Neurons in the prefrontal cortex have sophisticated projection patterns, referred to as *feedback* projections, which are optimally organized to allow the prefrontal cortex to modulate the activity of target regions (Harris et al., 2019; Gao et al., 2022). In particular, prefrontal feedback modulation of downstream regions is considered a key component of diverse cognitive operations, modulating the activity of cortical regions to regulate sensory-motor information and the activity of subcortical regions to regulate internal limbic drive (Miller, 2001; Kamigaki, 2018; Merre et al., 2021; Kenwood et al., 2022). The prefrontal cortex is thus an immediate region of interest in RTT due to its potential involvement in multiple cognitive, sensory-motor and emotional phenotypes. Previous work has revealed reduced activity of the prefrontal cortex in Mecp2 mutant mice (Kron et al., 2012). In addition, the discrete removal of MeCP2 in the forebrain replicates many behavioral phenotypes observed in MeCP2 knockout mice, including impaired motor coordination and increased anxiety (Gemelli et al., 2006). However, previous work has not addressed the potential impact of RTT on the prefrontal cortex’s ability to modulate the activity of downstream regions or analyzed structural alterations to the extensive network of prefrontal cortex feedback projections to cortical and subcortical structures.

Environmentally enriched housing is a behavioral intervention to stimulate motor and cognitive behaviors (van Praag et al., 2000), which has also been shown to alleviate some symptoms of *Mecp2* mutant mice. In particular, early environmental enrichment rescues motor coordination deficits (Kondo et al., 2008), and reduces anxiety behaviors in RTT mouse models (Lonetti et al., 2010). Enriched housing also results in increased synaptogenesis and increased levels of neurotrophic factors in both *Mecp2* mutant mice (Kondo et al., 2008; Lonetti et al., 2010) and wild-type mice (Baroncelli et al., 2010). While the majority of these changes have been reported in the hippocampus and cerebellum in RTT model mice, the impact of environmental enrichment on the prefrontal cortex remains unknown. In wild-type mice, enrichment induces an increase in synaptic density and dendritic arborization of prefrontal cortex neurons (Smail et al., 2020). Thus, environmentally enriched housing can produce diverse behavioral and synaptic changes in both *Mecp2* mutant and wild-type mice. An outstanding question is whether behavioral changes observed in RTT mouse models can be related to neuroanatomical alteration of prefrontal cortex projections, and whether these projections are affected by environmental enrichment.

Here, we carefully mapped the long-range axonal connections of the Anterior Cingulate Cortex (ACA), a prefrontal cortex region with extensive projections across the cortex. This allowed us to comprehensively evaluate if the reduction of MeCP2 in heterozygous female *Mecp2^+/−^*mice (the closest mouse model for female RTT patients) influences the neuroanatomical outputs of this key subdivision of the prefrontal cortex. In addition, we evaluated if enriched housing altered the projection pattern of the ACA, and how this related to expression levels of BDNF and behavioral phenotypes of *Mecp2^+/−^* mice. By comparing *Mecp2^+/−^*mice raised in either standard or enriched housing with wild-type controls, we found that ACA connectivity with the motor and somatosensory cortex was altered in *Mecp2^+/−^* female mice. Enriched housing reversed motor and anxiety phenotypes of *Mecp2^+/−^*mice, rescued BDNF levels in the hippocampus but not the prefrontal cortex, and surprisingly did not affect ACA projections to cortical or subcortical targets. Our data thus support the behavioral benefits of early environmental enrichment and suggest that these benefits can occur without large-scale changes in connectivity in Rett-model mice.

## MATERIAL AND METHODS

### Animals

All experiments were conducted using female mice, including both wild-type C57BL/6J (Jackson stock no. 000664) and transgenic heterozygous *Mecp2* knockout mice (*Mecp2^+/−^*) from the strain B6.129P2(C)-Mecp2^tm1.1Bird/J (Jackson stock no. 003890). Upon arrival at our facility at 4 weeks of age, all mice were randomly assigned to their respective housing conditions and immediately had small ear clip samples taken to confirm their genotypes. All experimental procedures and housing conditions adhered strictly to NIH guidelines and were approved by the MIT Animal Care and Use Committee.

### Housing

#### Feeding and Lighting Conditions

Both the enriched and standard housing groups were provided with food and water *ad libitum*. The housing facility maintained a reverse 12-hour light-dark cycle, with lights off during the daytime and on at night. This schedule was designed to align with the nocturnal activities of mice, thereby supporting their natural circadian rhythms and behavioral patterns.

#### Standard Housing Condition

Mice assigned to the standard environment were housed in Ancare N10 plastic rodent cages (7½” W x 11½” L x 5” H). These cages were equipped solely with nesting material to provide a basic environment without additional enrichment stimuli. The mice were housed communally in groups of four to five littermates per cage.

#### Enriched Housing Condition

Mice selected for the enrichment were placed in large Ancare N40 plastic rodent cages (10½” W x 19” L x 6⅛” H) at 4 weeks old. Each cage housed ten littermates communally. The enriched environment was furnished with a variety of certified enrichment toys provided by Bio-Serv™, including Mouse Igloos, Mouse Tunnels, Mouse Huts, Fast-Trac Activity Wheels, and Pup Tents, supplemented with rubber ball toys and bell toys. Enrichment objects were strategically rearranged every two days and completely swapped out weekly to sustain environmental novelty. The enrichment conditions commenced when the mice were 4 weeks of age and continued throughout the duration of the study.

### Experimental groups

As part of the experimental design, 5 out of the 10 mice in each experimental group were randomly selected to undergo intracranial viral injection surgeries aimed at quantifying axonal projections. Mice housed in the enriched environment were relocated before surgery to a separate enriched cage of identical size and equipped with the same assortment of toys and enrichment factors. Mice that underwent intracranial injections were given a 4-week period post-injection to allow for adequate viral expression. Following this period, they were euthanized and perfused at 12 weeks of age to facilitate detailed anatomical analyses.

Mice that did not receive intracranial injections were utilized for behavioral assessments after four weeks of standard or enriched housing. Following the conclusion of the behavioral tests, these mice were also euthanized, and their brains were carefully extracted and processed. Tissue samples were biopsied and preserved for subsequent quantification of brain-derived neurotrophic factor (BDNF) levels, which are indicative of neural health and plasticity. This synchronization ensured that data collected, whether anatomical, behavioral, or molecular, could be directly compared across different experimental conditions without age-related variability.

### Behavioral tests

Animals were held in a reverse dark-light cycle room. Behavioral experiments were conducted between 08:00 and 18:00, during the dark cycle. The testing environment was illuminated with ambient, red-tinted light, directed towards the walls to minimize direct light exposure and mimic nocturnal conditions for the mice.

#### Rotarod Performance Test

The motor coordination and learning abilities of mice were assessed using an accelerating rotarod apparatus. The rotarod accelerated from 4 revolutions per minute (rpm) to 40 rpm over a duration of 300 seconds. Each subject was evaluated on the rotarod for three trials separated by at least 10 minutes of rest in their home cage over the course of three consecutive days. The mean latency to fall per day was used for statistical analysis. The experimenter was blind to the genotype and housing condition of the mice being assessed.

#### Open Field Assay

The open field assay was conducted to evaluate locomotor activity and anxiety-like phenotypes. Briefly, animals were placed in the center of a 63.5 cm square plexiglass enclosure with bright illumination in the center of the field. Movements of the mice were recorded from an overhead perspective using a Panasonic Lumix DMC-FZ200 12.1 MP Digital Camera, set to capture video at a resolution of 1080p and a frame rate of 30Hz. All animals were tested without investigators in the room.

Animal tracking analysis was performed using the ‘Detect Any Mouse Model’ (DAMM) software, developed by Kaul et al. (Kaul et al., 2024). This software employed a pre-trained object detection algorithm optimized for localizing mice within complex environments. To quantify the locomotor activity, the x and y coordinates of the mouse in each frame were used to calculate distance traveled, velocity and percentage of time spent in the center quadrant and corners of the arena during the first 5 minutes in the arena. The center region of the arena was defined by shrinking the border rectangle toward its centroid by a factor of 0.7. The corners of the arena were defined by creating polygons around each corner point, extending lines toward adjacent corner points by a factor of 0.3.

### Anatomical analysis

#### Viral injections

Mice were anesthetized with isoflurane (1.5%) and given preemptive analgesia (extended-release Buprenorphine, 1mg/kg,). After hair removal and sterilization of the skin with 70% ethanol and iodine each animal was placed into a stereotaxic frame (51725D, Stoelting). Body temperature was maintained at 36°C with a feedback-controlled heating pad (ATC2000, World Precision Instruments). For viral injections, a micropipette attached on a Quintessential Stereotaxic Injector (QSI 53311, Stoelting) was used. A small craniotomy was drilled above the right Anterior Cingulate Cortex (AP: +1, ML: −0.3, DV: −0.9) and 0.2 µl of AAV1-CAG-tdTomato (59462-AAV1, Addgene, 5×10¹² vg/mL) was injected unilaterally at a rate of 0. 05µl/min. Virus expression was allowed for 4 weeks before the mouse was perfused for tissue collection. The pipette was held in place for 5 min after each viral injection before being slowly retracted from the brain. Postoperative analgesics (meloxicam 5mg/kg) was given at the end of the surgery, followed by a second dose 18-24h after the surgery, and recovery was monitored for a minimum of 72 h after surgery.

#### Perfusion and sectioning

Following an overdose of isoflurane, all mice were transcardially perfused with 1x phosphate buffer (PBS) followed by 4% paraformaldehyde (PFA) in PBS. Brains were left overnight in 4% PFA and washed three times in PBS the following day. For sectioning, perfused brains were embedded in 4% UltraPure™ Agarose for stabilization and glued to the specimen disk of a vibrating microtome (Leica VT1200S). 80 µm thick tissue sections were collected sequentially from the most anterior part of the cortex, immediately posterior to the olfactory bulb, to the most posterior part of the cortex, just anterior to the cerebellum. The collected sections were placed in 24-well plates filled with PBS and stored at 4°C.

#### Brain section preparation and staining

Every other section was collected and placed free-floating for incubation with Hoechst staining (Thermo Scientific™ Hoechst 33342 solution, 1:2,000 in PBS) for 5-10 minutes. The sections were thereafter washed in PBS three times and mounted onto microscope slides (VWR® Premium Superfrost® Plus Microscope Slides, 48311-703). Finally, each slide was covered with a coverslip using VECTASHIELD® HardSet™ Antifade Mounting Medium (Fisher Scientific, NC9029229).

#### Image acquisition

Brain sections were imaged using an epifluorescence slide scanning microscope (Zeiss, Tissue Gnostics TissueFAX Slide Scanner) equipped with a 10x, 0.45 NA acquisition air objective, Chroma multiband filter sets allowing for fast scanning of multiple colors, and a sCMOS camera (Hamamatsu C15440-20UP). Automated tissue detection with parameters targeting sections greater than 2 mm² or 12,006 pixels was used. Imaging illumination parameters were optimized with tissue from several animals, and all tissue sections were acquired using the same settings. Detailed images of each brain section were captured in two wavelengths corresponding to Hoechst staining (blue, 460nm) and tdTomato (red, 580nm). Each 80µm section of brain tissue was imaged at 5 focal planes 10µm apart, covering 50µm distance through the volume. The maximum projection of these sections through the volume were used for quantification. Automated image stitching was used to compile seamless composites of each brain section.

#### Image correction and enhancement

Due to the relative brightness near the injection site, discrete axonal projections further away were comparatively dim and difficult for the automated axon detection pipeline to detect. To ensure consistent and accurate detection of axon projections throughout the entire brain, we implemented a custom gamma adjustment protocol using a Fiji macro. Briefly, the average intensity value across all images was calculated by summing the mean intensity values for all brain slice images and dividing this total by the number of images in the dataset for each mouse. This average intensity value served as a reference baseline for subsequent fluorescent intensity adjustments. For the most anterior quarter of images, the gamma correction value for each image was calculated based on the ratio of the image’s mean intensity to the overall average intensity of all images in the dataset. The ratio was constrained to ensure that it did not fall below a lower limit of 1.0 (no correction) or exceed an upper limit of 1.5. This approach allowed for gamma reductions for images with higher mean intensities compared to the overall average but prevented excessive dimming by limiting the correction range.

For the last quarter of images, i.e., the most posterior sections, the gamma value was calculated in a similar manner by dividing each image’s mean intensity by the overall average intensity. However, since these images were generally dimmer, the gamma correction was adjusted with a lower bound of 0.9 to slightly enhance the fluorescent intensity. The upper limit for the correction was capped at 1.0 to prevent over-enhancement. This ensured that the images were brightened just enough to compensate for the reduced signal strength, while preserving the overall structure and fidelity of the fluorescence data. This differential approach accounted for potential variations in signal strength across the image series. Images of brain sections with bregma coordinates between the most anterior and posterior quarters where not adjusted.

#### Whole brain reconstruction and quantification

To quantitatively compare projection densities we utilized the BrainJ pipeline (Botta et al., 2020) to reconstruct tissue sections into whole brains and aligned them to the Allen Brain Reference Atlas to allow for quantitative comparisons of projection density. For registration and alignment, the Hoechst channel was used. In order to analyze the target channel containing our axons, several settings in BrainJ were implemented to optimize the detection of axonal projections. The reference section for initial registration was typically chosen from the middle of the dataset to serve as a baseline for aligning other sections. Background noise was reduced using a rolling ball filter with a radius of 2, suitable for a resolution of 2 μm per pixel. The analysis of projections was conducted using the Binary Threshold method, which detected any fluorescence above a set threshold in the mesoscale mapping analysis. The set threshold was adjusted for individual sections to prevent oversaturation and oversampling around the injection site.

#### Axon density analysis

In our analysis of axonal projections, we employed a custom Python script to preprocess the data obtained from BrainJ. For analysis and visualization of the data we used the Allen Brain Atlas nomenclature and structure tree as a reference. Each target region was also confirmed by comparison to the Allen Brain Reference Atlas connectome database.

The BrainJ analysis pipeline produced two measurements of axon density for each brain region: ‘axon density’ and ‘relative density’. ‘Axon density’ was defined as the total volume of ‘axon pixels’ in a region divided by the total volume of pixels of the region. ‘Relative density’, expressed as a percentage, was defined as the total volume of ‘axon pixels’ in a specific brain region divided by the total volume of ‘axon pixels’ detected across the entire brain. The ‘relative density’ value accounted for differences in the efficiency of virus uptake or transport and was used for further analysis. We also visualized the data as ‘proportional density’, defined as the relative density in one area (for example, VISp) normalized by the relative density in a defined part of the brain (eg., the cortex). The proportional density yielded a comparison between brain regions or subregions independent of the overall efficiency of viral labeling in each brain. The proportional density was used for conclusions regarding the distribution of axons in a given region across various experimental conditions, and relative density was used to describe differences in actual projection density.

### Total protein quantification and BDNF ELISA

Upon completion of behavioral testing, mice were euthanized and brain tissue was collected for the measurement of brain-derived neurotrophic factor (BDNF) protein levels. Mice were decapitated, the whole brain was rapidly removed and the prefrontal cortex of both hemispheres were dissected out using an Integra Miltex Disposable 3 mm biopsy punch. Additionally, the entire hippocampus from both hemispheres were manually microdissected. All tissue was flash frozen on dry ice and stored at −80C till further use. The frozen tissue samples were lysed in 200uL of lysis buffer containing 150mM NaCl, 1% NP-40, 50mM Tris-HCl (pH 7.5), 5mM EDTA, and 1X protease cocktail (Cell Signaling). The samples were repeatedly passed through a sterile 25G needle and 1mL syringe until fully lysed and homogenized. The resulting lysate was centrifuged for 10 min at 14,000 g and the supernatant was used for estimation of BDNF and protein concentrations. BDNF levels were quantified using the Total BDNF ELISA Kit (Quantikine, bio-techne, R&D Systems) according to the manufacturer’s instructions. Total protein concentration was quantified using the Pierce™ Bradford Plus Protein Assay Kit (Thermo Scientific) according to the manufacturer’s instructions. BDNF levels for each sample were normalized to the total protein concentration of that sample.

### Statistical analysis

Statistical analysis was performed in GraphPad Prism version 10.0.0, or in Python 3.8.8 using the sklearn and scipy libraries. Details of all statistical comparisons are noted in the text.

## RESULTS

### *Mecp2^+/−^* mice show behavioral deficits that can be rescued with enriched housing

As a first step in characterizing the involvement of the ACA in the behavioral deficits seen in *Mecp2^+/−^* female mice, we measured locomotion, motor coordination and anxiety (Stearns et al., 2007) using the rotarod and open field assays. Three experimental groups were compared: wild type female mice housed in standard housing (WT), *Mecp2^+/−^* female mice housed in standard housing (*Mecp2^+/−^* SH), and *Mecp2^+/−^* female mice that had four weeks of enriched housing prior to behavioral testing (*Mecp2^+/−^*EH) (**Figure 1a-b**). All mice were 8 weeks old at the time of behavioral testing.

**Figure 1.**
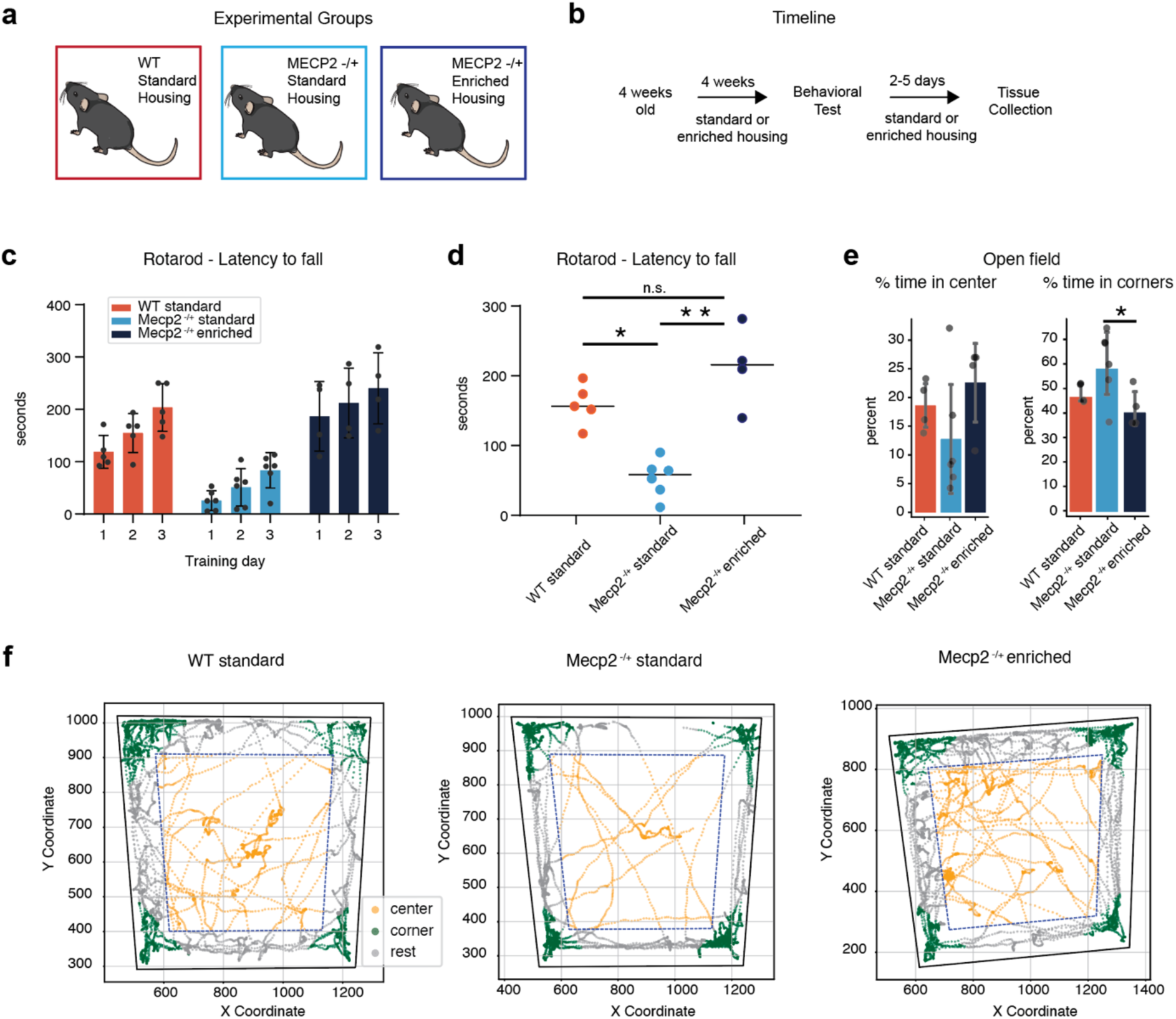
Open Field Assay shows motor deficit phenotype and anxiety-like behavior in Rett mice. **(a)** Schematic of experimental groups. **(b)** Experimental timeline. **(c)** Mean latency to fall off the rotarod for three consecutive test days for WT standard housing (red, n=5 mice), Mecp2 standard housing (light blue, n=6 mice) and Mecp2 enriched housing (blue, n=4 mice) experimental groups. Bars represent mean values, error bars are standard error and points represent individual animals. **(d)** Mean latency to fall across all test days for WT standard housing (red, n=5 mice), Mecp2 standard housing (light blue, n=6 mice) and Mecp2 enriched housing (blue, n=4 mice) experimental groups. Lines represent mean values and points represent individual animals. *p-value < 0.05, **p-value < 0.0001, one-way ANOVA and Tukey’s post-hoc test. p-value = Mecp2 SH-Mecp2 EH:<0.0001, Mecp2 SH-WT: 0.016. **(e)** Percent of time spent in the center and corners of the open field of WT standard housing (red, n=5 mice), Mecp2 standard housing (light blue, n=6 mice) and Mecp2 enriched housing (blue, n=4 mice) experimental groups. Bars represent mean values; error bars are standard error and points represent individual animals. *p-value < 0.05, one-way ANOVA and Tukey’s post-hoc test. p-value = Mecp2 SH-Mecp2 EH: 0.047. **(f)** Example trace of a mouse moving in the open field, in the center (orange), corners (green) or rest (grey) of the arena, for each experimental group.

*Mecp2^+/−^* SH mice showed impaired motor coordination in the rotarod assay and had a significantly shorter average latency to fall compared to WT mice (**Figure 1c-d**; animal numbers and statistical tests are noted in the figure legend). The observed impairment was fully rescued with environmental enrichment and the *Mecp2^+/−^* EH group displayed no difference in average latency to fall compared to the WT group (**Figure 1d**). Although the *Mecp2^+/−^* SH did considerably worse in the rotarod assay than the WT and *Mecp2^+/−^* EH group, all mice increased their latency to fall on each subsequent day of testing, indicating a sequential learning of the task by all groups (**Figure 1c**).

Loss of MeCP2 either brain-wide (Lonetti et al., 2010), in the forebrain (Gemelli et al., 2006) or in the amygdala (Adachi et al., 2009) induces anxiety in mice as measured by thigmotaxis in an open field arena. We evaluated if enriched housing of *Mecp2^+/−^*mice could reduce such anxiety. *Mecp2^+/−^* SH mice showed a trend toward spending less time in the center of the arena on average compared to the other two groups, and spent significantly more time in the corners relative to the *Mecp2^+/−^* EH group (**Figure 1e-f**). The *Mecp2^+/−^* EH group did indeed show reduced anxiety and spent more time in the center than both the *Mecp2^+/−^* SH and WT groups (**Figure 1e-f**). We did not observe a motor deficit of the gait of the *Mecp2^+/−^* SH group in comparison to the WT group. Instead, *Mecp2^+/−^* EH mice had a higher average velocity and longer cumulative distance traveled in the open field than the other two experimental groups (**Supplementary Figure 1**).

Our results thus confirm that enriched housing rescues the fine motor skill deficit seen in *Mecp2^+/−^*female mice, with *Mecp2^+/−^* EH mice performing the rotarod assay comparably to the WT group (**Figure 1c-d**) (Kondo et al., 2008; Lonetti et al., 2010). In addition, enriched housing also reduced the anxiety phenotype of *Mecp2^+/−^*mice (**Figure 1e-f**).

### Comparing whole-brain projection density of forebrain axons in wild-type and *Mecp2^+/−^* mice

The anxiety phenotype of *Mecp2^+/−^* mice has previously been attributed to the loss of MeCP2 protein in the forebrain(Gemelli et al., 2006). In addition, the frontal cortex has reduced activity in *Mecp2^+/−^*mice (Kron et al., 2012) and reduced size in RTT-patients (Takeguchi et al., 2022). Therefore, following the results of our behavioral testing, we characterized the anatomical patterns of PFC connectivity in *Mecp2^+/−^* mice, and potential structural changes as a consequence of enriched housing. To carefully map the axon density of ACA projection neurons on a brain-wide scale, AAV-CAG-tdTomato was injected into the ACA of WT, *Mecp2^+/−^* SH and *Mecp2^+/−^* EH mice (**Figure 2a-b**). Each experimental group had experienced four weeks of enriched or standard housing prior to the viral injection and continued in the same housing condition for an additional four weeks, after which the mice were perfused and the tissue collected (**Figure 2c**). The whole brain was coronally sectioned, stained with Hoechst stain, and imaged (**Figure 2d**). The axonal density of ACA axons in each brain region was thereafter quantified following three steps within the ImageJ plugin BrainJ (Botta et al., 2020). First, the location of ACA axons in each image was saved as a mask by the segmentation of each axon-filled pixel based on fluorescent intensity in the red channel. Second, each image was aligned to the Allen Brain Reference Atlas (Lein et al., 2007) based on landmarks in the Hoechst channel. Third, the detected axon pixels were overlaid with the Allen Brain Reference Atlas (**Figure 2e**). A data file was generated for each brain, containing an ‘axon density’ and ‘relative density’ value for each brain region within the Allen Brain Reference Atlas. (see Methods *‘Axon density analysis’* for details; complete list of abbreviations in **Supplementary Table 1**).

**Figure 2.**
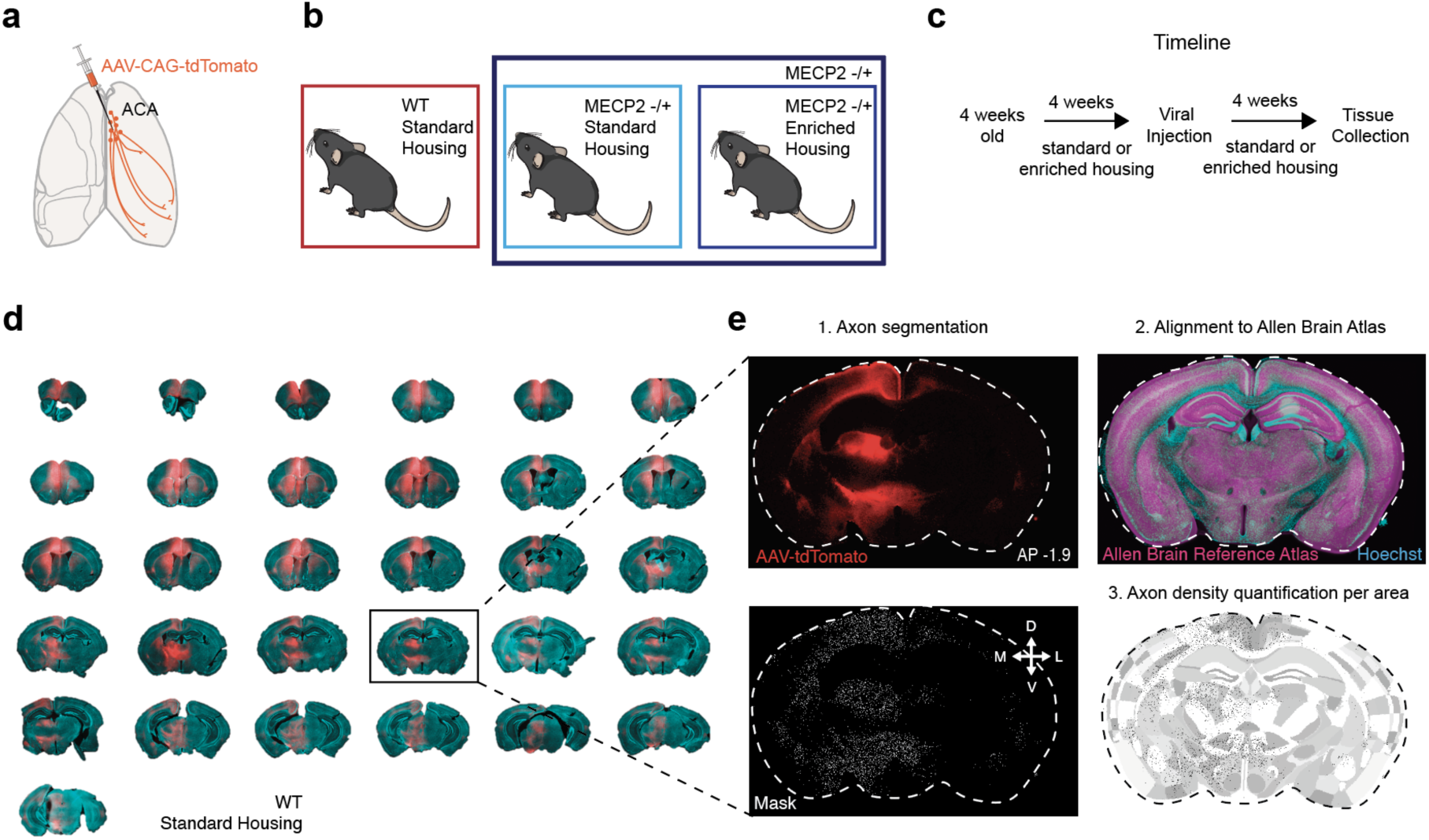
Pipeline for tracing studies and anatomical data collection and analysis. **(a)** Schematic of experimental strategy. AAV-CAG-tdTomato was injected into the ACA of Mecp2 and WT mice. **(b)** Three experimental groups were compared, WT mice in standard housing, Mecp2 in standard housing and Mecp2 in enriched housing. **(c)** Experimental timeline. **(d)** Coronal brain sections of WT mouse with td-Tomato expression in ACA axons (red) and Hoechst stain (cyan) staining for whole-brain processing of ACA axon density. **(e)** Each brain section was processed in three steps. 1. ACA axons (up) were segmented out and each pixel of axon saved as a mask (down). 2. Each section was aligned to the Allen Brain Reference Atlas (ARA) using the Hoechst stain channel (cyan). 3. Segmented axons were then aligned to the ARA and a density measurement was calculated based on the number of pixels of axons within each region divided by the number of pixels of the area of the region.

### *Mecp2^+/−^* mice show altered connectivity to somatosensory and motor regions that are unaffected by enriched housing

The ACA has extensive connectivity to the rest of the cortex, including medial, lateral, and posterior regions (Zingg et al., 2014; Ährlund-Richter et al., 2024). As described previously in WT mice, ACA axons also innervate the majority of cortical regions in *Mecp2^+/−^* mice (**Figure 3a, Supplementary Table 1**). Comparing the ACA axon density in each cortical region of the ipsilateral hemisphere, we observed no difference between the *Mecp2^+/−^* SH and *Mecp2^+/−^* EH experimental groups (**Figure 3a, left**). However, when we compared the axon density of ACA projections in all *Mecp2^+/−^* mice, regardless of housing conditions, to WT mice we found differences in key cortical regions. *Mecp2^+/−^*mice had a significantly larger proportion of ACA axons within the somatosensory cortex (SS) in comparison to WT mice (**Figure 3a, right, Supplementary Figure 2**). Conversely, the proportional density of ACA axons in the motor cortex (MO) was decreased in the *Mecp2^+/−^*mice in comparison to WT mice (**Figure 3a, right**).

**Figure 3.**
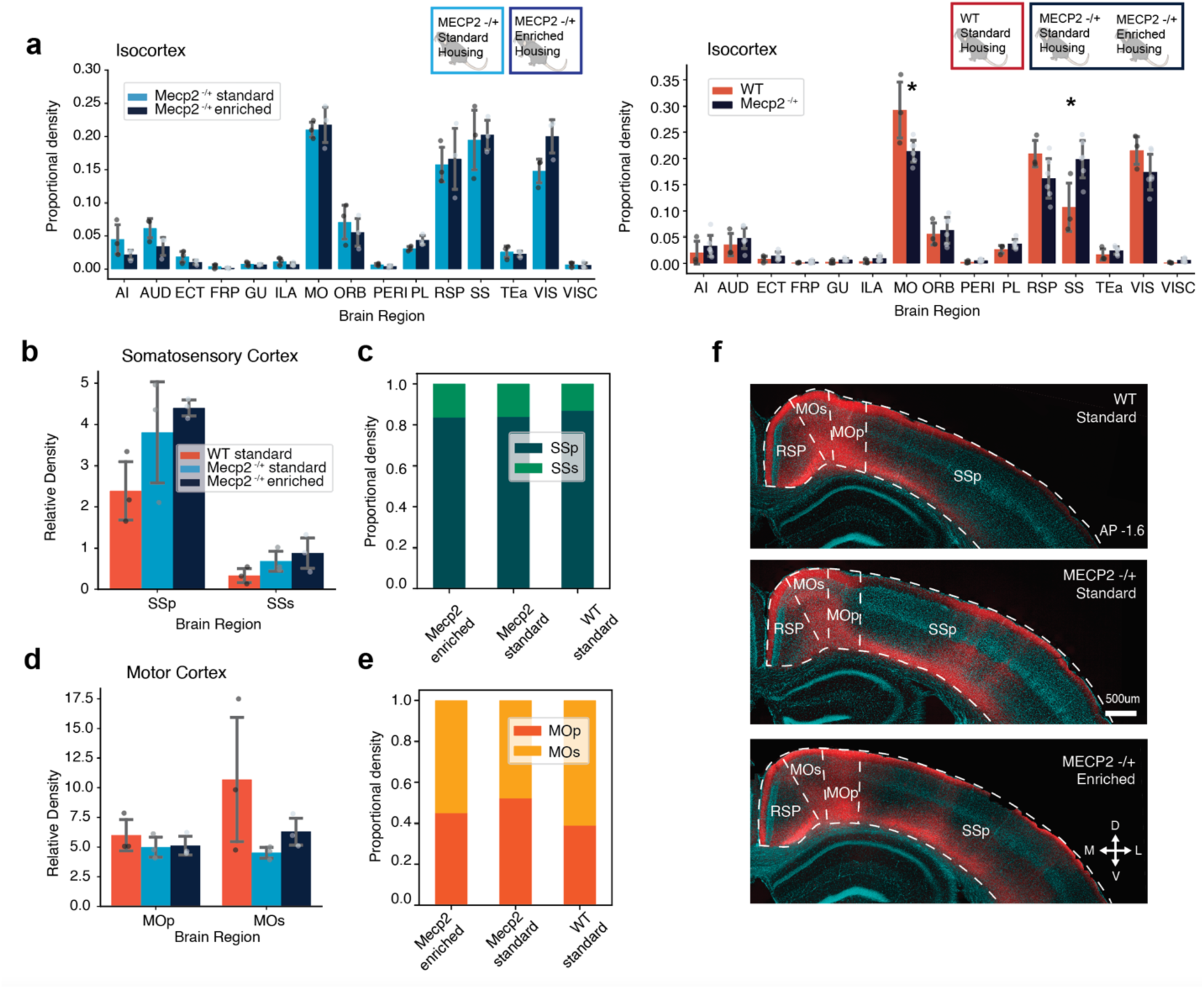
ACA axonal projections in *Mecp2^+/−^* mice are denser in the somatosensory cortex and sparser in the motor cortex in comparison to wild-type mice. **(a)** (Left) Proportional density of ACA axons per cortical region out of all of Isocortex in Mecp2 standard housing (light blue, n=3 mice) and Mecp2 enriched housing (dark blue, n=3 mice) mice. (Right) Proportion of Relative Density of ACA axons per cortical region out of all Isocortex comparing WT standard housing (red, n=3 mice), Mecp2 (blue, n=6 mice) mice. *p-value < 0.05, unpaired Student’s t-test. MO p-value: 0.027, SS p-value: 0.023. **(b)** Relative Density of ACA axons in SS in WT standard housing (red, n=3 mice), Mecp2 standard housing (light blue, n=3 mice) and Mecp2 enriched housing (blue, n=3 mice) mice. **(c)** Proportional density of ACA axons in SSp (dark green) vs. SSs (green) comparing WT standard housing (n=3 mice), Mecp2 standard housing (n=3 mice) and Mecp2 enriched housing (n=3 mice) experimental groups. **(d)** Relative Density of ACA axons in MO WT standard housing (red, n=3 mice), Mecp2 standard housing (light blue, n=3 mice) and Mecp2 enriched housing (blue, n=3 mice) mice. **(e)** Proportional density of ACA axons in MOp (orange) vs. MOs (yellow) comparing WT standard housing (n=3 mice), Mecp2 standard housing (n=3 mice) and Mecp2 enriched housing (n=3 mice) experimental groups. **(f)** Representative images of ACA axons (red) in RSP, MO and SS in WT (up), Mecp2 standard housing (middle) and Mecp2 enriched housing (down) mice. Complete list of abbreviations in **Supplementary Table 1.**

The increased proportion of ACA axon density in the somatosensory cortex of *Mecp2^+/−^* mice was the result of increased density of axons in both the primary and secondary somatosensory cortex, for both the standard and enriched housing experimental groups (**Figure 3b, f**). The proportional density of ACA axons between the primary and secondary somatosensory cortex was comparable between WT, *Mecp2^+/−^* SH and *Mecp2^+/−^* EH experimental groups (**Figure 3c**). WT mice showed a slight increase in ACA relative axon density in the secondary motor cortex in comparison to the primary motor cortex, which *Mecp2^+/−^* mice did not (**Figure 3d**). ACA axons were evenly distributed between the primary and secondary motor cortex in *Mecp2^+/−^* SH and *Mecp2^+/−^*EH mice, with no significant differences observed (**Figure 3e**).

Overall, our results show that the relative ACA axon density is increased in the somatosensory cortex of *Mecp2^+/−^* mice in comparison to WT mice and is due to an increase in ACA axon density in both the primary and secondary somatosensory cortex. The observed decrease in the proportion of ACA axon in the motor cortex of *Mecp2^+/−^* mice (**Figure 3a, right)** appears to be in part due to a decrease in relative axon density in the secondary motor cortex (**Figure 3d**). Neither of the anatomical structural changes observed in the *Mecp2^+/−^*mouse model were affected by enriched housing.

### *Mecp2^+/−^* mice show altered distribution of ACA axons within the major cortical projection profiles of PFC axons

Recent large-scale anatomical mapping of prefrontal cortex projection neurons has revealed four major projection patterns of prefrontal cortex output neurons, referred to as *PFC networks*(Gao et al., 2022). These projection profiles sort projection neurons as targeting either the prefrontal cortex, the central cortex, the medial cortex, or the lateral cortex (**Figure 4a**). To further characterize structural changes of the ACA in *Mecp2^+/−^* mice we examined the axonal density of ACA axons within each major PFC network. *Mecp2^+/−^* SH and *Mecp2^+/−^*EH experimental groups showed no difference in the distribution of ACA axons within each PFC network (**Figure 4b**). The largest proportion of ACA axon density in both *Mecp2^+/−^*groups was observed in the medial network (**Figure 4b**), which includes cortical regions such as the visual cortex and retrosplenial cortex. This aligns with previous reports that show the ACA is a major source of input to the visual and retrosplenial cortex in wild-type mice (Zingg et al., 2014; Zhang et al., 2016; Merre et al., 2021; Ahrlund-Richter et al., 2024; Liu et al., 2024b). Comparing the distribution of ACA axons within each PFC network in *Mecp2^+/−^* and WT mice we found that *Mecp2^+/−^* mice had a significantly higher proportion of ACA axons in the central network in comparison to WT mice (**Figure 4c**). The increase in ACA axons within the central network is in line with the observed increase in ACA axons in the somatosensory cortex (**Figure 3a-b and Supplementary Figure 3**), as the central network consists mainly of the somatosensory cortex and primary motor cortex. WT mice instead showed a slight increase in ACA axons within the medial and prefrontal network (**Figure 4c**).

**Figure 4.**
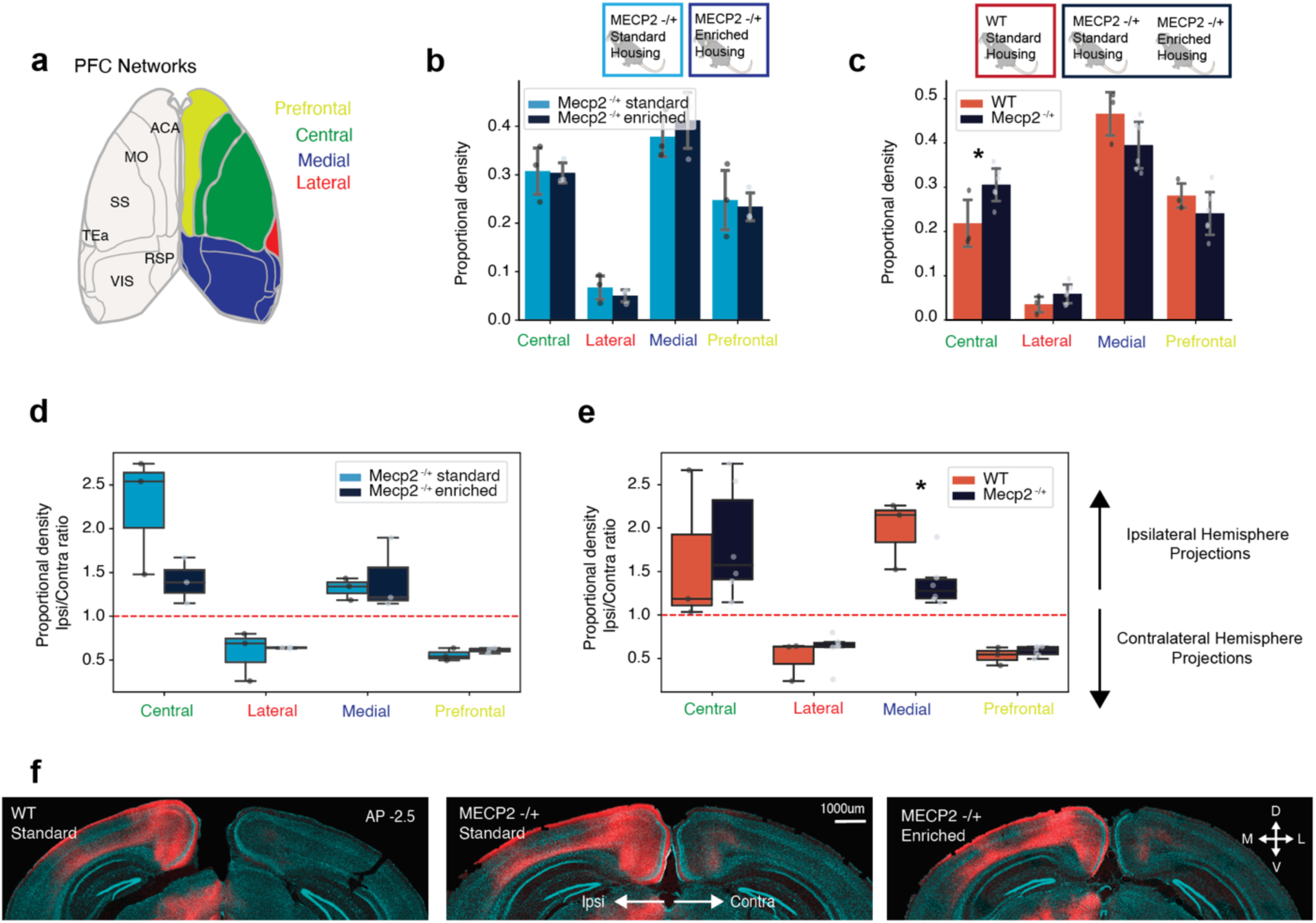
ACA axons more densely participate in the central PFC network in *Mecp2^+/−^* mice. **(a)** Schematic of cortical regions participating in each PFC network. Prefrontal: FRP, PL, ILA, ORB, ACA, MOs, AId, AIv. Lateral: AIp, GU, VISC, TEa, PERI, ECT, ENT. Central: MOp, SS. Medial: PTLp, RSP, VIS, AUD. **(b)** Proportional density of ACA axons for brain regions participating in each PFC network for Mecp2 standard housing (light blue, n=3 mice) and Mecp2 enriched housing (blue, n=3 mice) experimental groups. Bars represent mean values; error bars are standard deviation and points represent individual animals. **(c)** Proportional density of ACA axons for brain regions participating in each PFC network in WT (red, n=3 mice) and Mecp2 (dark blue, n=6 mice) mice. *p-value < 0.05, unpaired Student’s t-test. Central p-value: 0.039. **(d)** Relative Density proportion between ipsilateral and contralateral hemispheres for ACA axons innervating each PFC network for Mecp2 standard housing (light blue, n=3 mice) and Mecp2 enriched housing (blue, n=3 mice) experimental groups. **(e)** Relative Density proportion between ipsilateral and contralateral hemispheres for ACA axons innervating each PFC network in WT (red, n=3 mice) and Mecp2 (dark blue, n=6 mice) mice. *p-value < 0.05, unpaired Student’s t-test. Medial p-value: 0.030. Each boxplot represents the quartiles of values, while the whiskers extend to show the rest of the values within 1.5 times the interquartile range. Points represent individual animals. **(f)** Representative images of ACA axons (red) of the Medial Network (RSP, VIS) in WT standard housing (left) and Mecp2 standard housing (middle) and Mecp2 enriched housing (right) mice. Complete list of abbreviations in **Supplementary Table 1.**

The distribution of cortical axons between the ipsilateral and contralateral hemisphere is configured during development, and considered critical for the coordination of sensory-motor and cognitive functions (Zhou et al., 2013; Fenlon et al., 2017). ACA projection neurons can target either the ipsilateral or contralateral hemisphere, or both (Gao et al., 2022; Liu et al., 2024a). We observed that the absolute density of ACA axons was greater in the ipsilateral hemisphere for all regions, as expected (**Supplementary Figure 3d-e**). To investigate the potential shift in the distribution of ACA projections across the two hemispheres we compared the proportional ACA axon density for each PFC network as a ratio between the ipsilateral and contralateral hemisphere. We found that the ipsilateral hemisphere had a higher proportion of ACA axons in the central and medial network in all experimental groups (ratio above 1) (**Figure 4d-e**). There was no significant difference between the *Mecp2^+/−^* SH and *Mecp2^+/−^* EH groups in this distribution across hemispheres (**Figure 4d**). Comparing the WT group with the combined *Mecp2^+/−^* group we observed a significant difference in the proportional distribution of ACA axons across hemispheres in the medial network (**Figure 4e**): WT mice had a larger proportion of ACA axons within the medial network in the ipsilateral hemisphere in comparison to *Mecp2^+/−^* mice (**Figure 4f**).

Thus, our results indicate that ACA axons of *Mecp2^+/−^*mice participate to a larger degree in the central PFC network in comparison to WT mice. This results in a proportionally larger density of ACA axons in the medial PFC network in WT mice. Consequently, the proportional distribution of ACA axons across each hemisphere is redistributed in *Mecp2^+/−^*mice, and the ipsilateral projections are less dominant in the medial network in comparison to WT mice.

### *Mecp2^+/−^* mice display no difference in ACA subcortical projections

ACA neurons project to distinct subcortical target regions in addition to cortical structures. The role of the ACA in social and cognitive behaviors has been particularly investigated in relation to its connectivity with the thalamus and midbrain (Huda et al., 2020; Smith et al., 2021; Song et al., 2023). To evaluate if *Mecp2^+/−^* mice had altered ACA projection profiles in comparison to WT mice, and if this was affected by housing conditions, we compared the proportional distribution of ACA axon density in 8 major brain regions, including isocortex and 7 subcortical regions, as defined by the Allen Brain Reference Atlas (**Figure 5a**). We observed no difference in distribution of ACA axons, and all experimental groups showed a majority of ACA axons within the isocortex, followed by the striatum, midbrain, and thalamus (**Figure 5b**). ACA projections to the thalamus of WT, *Mecp2^+/−^* SH, and *Mecp2^+/−^*EH mice all showed previously documented connectivity with nuclei such as the mediodorsal thalamic nuclei, lateral pulvinar nuclei, posterior complex of the thalamus and ventral medial nucleus of the thalamus (Hunnicutt et al., 2014; Oh et al., 2014) (**Figure 5c and Supplementary Figure 4a**). The major target of ACA projections in all experimental groups in the striatum was the caudate putamen (**Figure 5d**), in particular the mediodorsal caudate putamen (Hunnicutt et al., 2018) (**as seen in Figure 2d**). Finally, ACA axons of WT, *Mecp2^+/−^* SH and *Mecp2^+/−^* EH targeted previously documented midbrain structures such as the superior colliculus and ventral tegmental area to an equivalent extent (**Figure 5e**).

**Figure 5.**
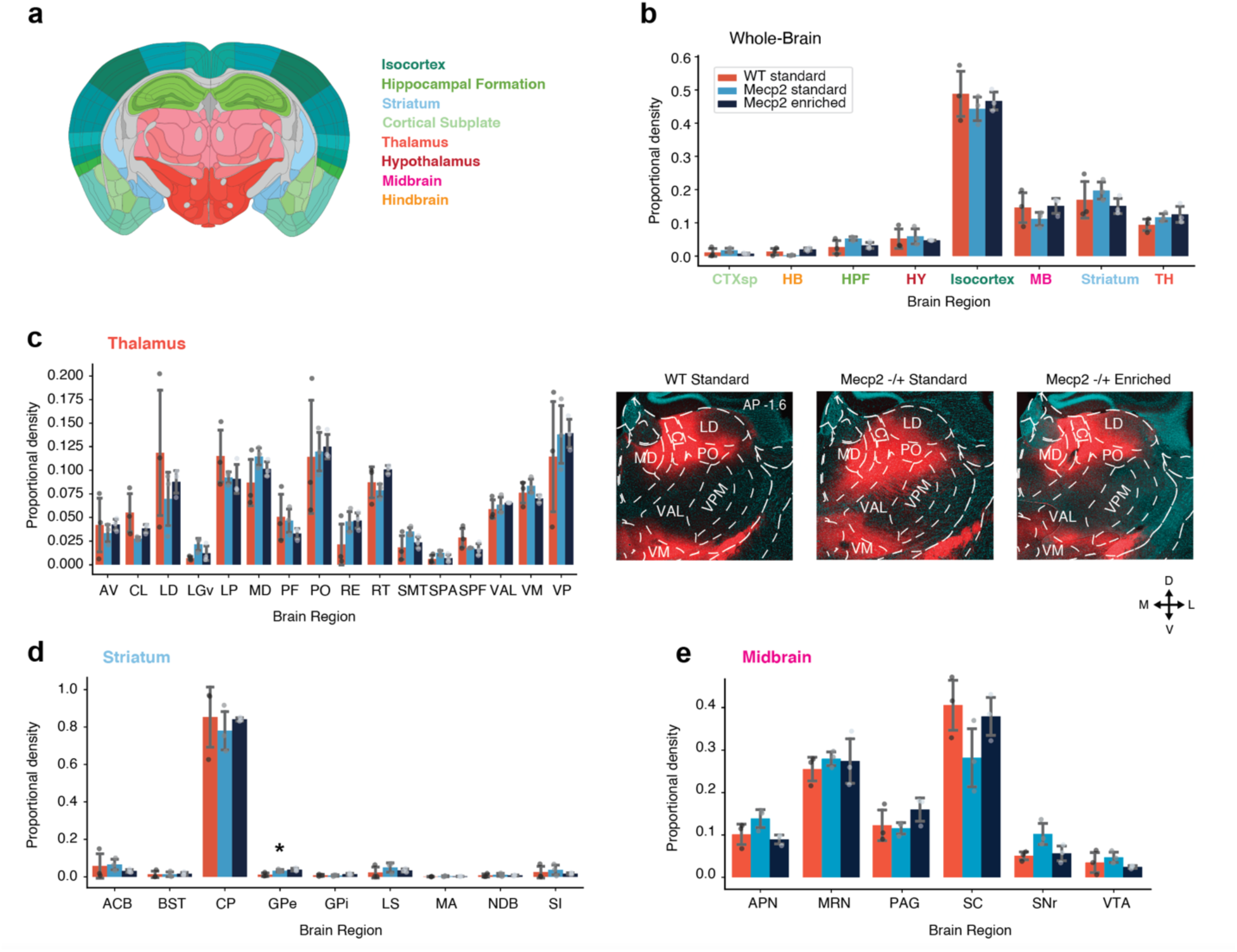
ACA axon density in subcortical structures. **(a)** Schematic representation of a coronal brain section from the Allen Brain Reference Atlas. Major brain regions listed and color coded as in the Allen Brain Reference Atlas. **(b)** Proportional density of ACA axons for each major brain region for WT standard housing (red, n=3 mice), Mecp2 standard housing (light blue, n=3 mice) and Mecp2 enriched housing (blue, n=3 mice) experimental groups. Abbreviations as in (a). **(c)** (Left) Proportional density of ACA axons in Thalamic nuclei for WT standard housing (red, n=3 mice), Mecp2 standard housing (light blue, n=3 mice) and Mecp2 enriched housing (blue, n=3 mice) experimental groups. Thalamic nuclei with a minimum of 1% of detected axonal density in each animal are included. Extended plot in Supplementary Figure 4A. (Right) Representative images of ACA axons (red) in Thalamus in WT standard housing (left) and Mecp2 standard housing (middle) and Mecp2 enriched housing (right) mice. **(d)** Proportional density of ACA axons in the Striatum for WT standard housing (red, n=3 mice), Mecp2 standard housing (light blue, n=3 mice) and Mecp2 enriched housing (blue, n=3 mice) experimental groups. **(e)** Proportional density of ACA axons in the Midbrain for WT standard housing (red, n=3 mice), Mecp2 standard housing (light blue, n=3 mice) and Mecp2 enriched housing (blue, n=3 mice) experimental groups. Bars represent mean values; error bars are standard deviation and points represent individual animals. Complete list of abbreviations in **Supplementary Table 1.**

Our results thus display a dichotomy of cortical and subcortical ACA projections of *Mecp2^+/−^* mice. While cortical ACA projections show specific differences in *Mecp2^+/−^* mice in comparison to WT mice, subcortical projections do not. Environmentally enriched housing of *Mecp2^+/−^* mice does not change the distribution or density of ACA axons in either cortical or subcortical structures.

### Enriched housing rescues BDNF levels in the hippocampus, but not prefrontal cortex, of *Mecp2^+/−^* mice

The neurotrophin BDNF influences synaptic plasticity and transmission and is known to be affected by Mecp2(Chen et al., 2003; Chang et al., 2006; Li and Pozzo-Miller, 2013); BDNF levels are decreased in *Mecp2^+/−^*mice and enriched housing rescues these reduced levels in the cerebellum and hippocampus (Kondo et al., 2008). Therefore, to examine the relationship between the rescue of behavioral deficits, BDNF levels, and ACA axonal projections, we measured the levels of BDNF in the prefrontal cortex of WT, *Mecp2^+/−^* SH and *Mecp2^+/−^* EH mice. The hippocampus was used as a control region and did indeed show decreased levels of BDNF in *Mecp2^+/−^* SH mice in comparison to the WT group (**Figure 6a**). The *Mecp2^+/−^* EH group showed a complete rescue of BDNF level in the hippocampus and was not significantly different from the WT group (**Figure 6a**). Intriguingly, the BDNF level in prefrontal cortex of the *Mecp2^+/−^* SH group was decreased to a similar extent as in the hippocampus, but was not rescued by the enriched housing condition of the *Mecp2^+/−^* EH group (**Figure 6b**). Our results therefore indicate that enriched housing reverses the motor and behavioral deficits of *Mecp2^+/−^* mice, as well as restores their hippocampal BDNF level, but does not change the level of BDNF in the prefrontal cortex.

**Figure 6.**
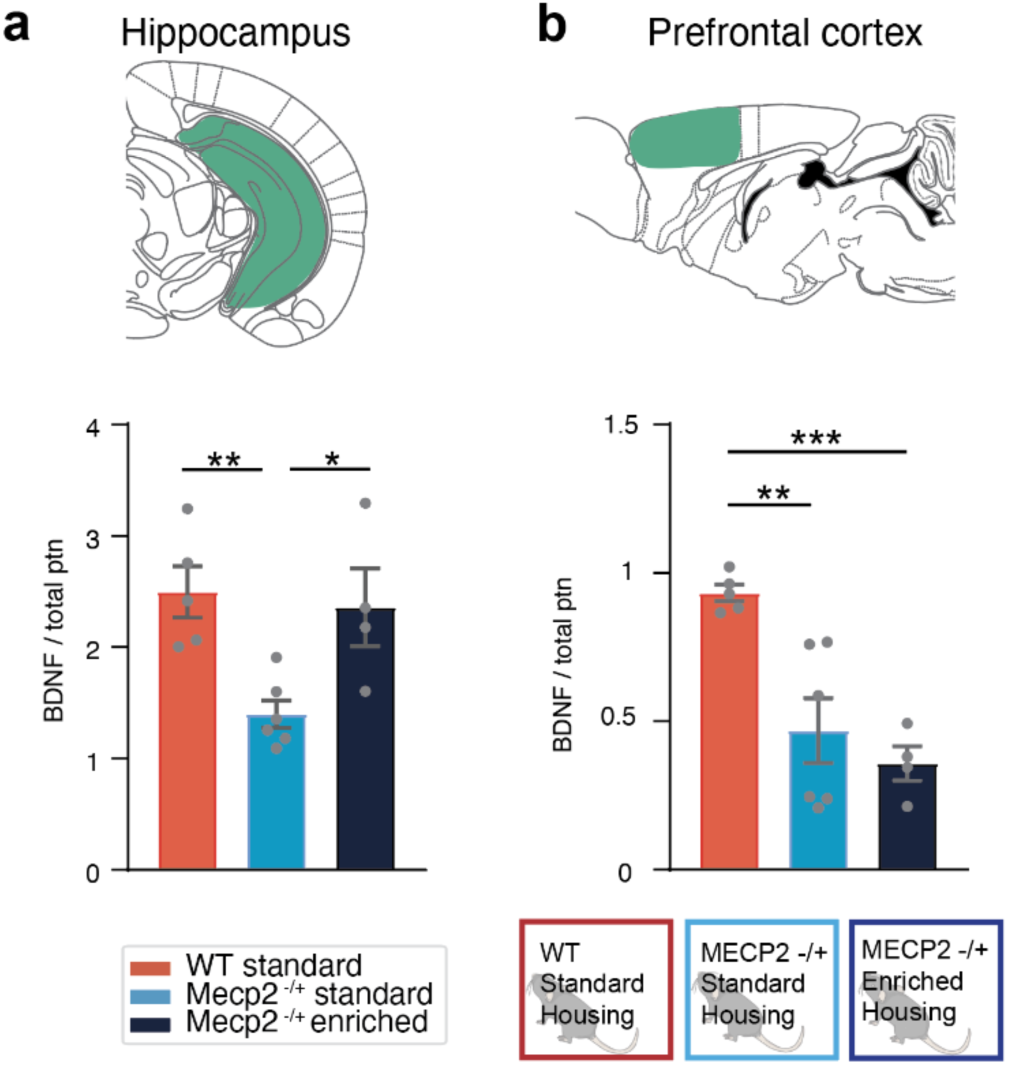
BDNF expression is decreased in RTT mice and restored in the hippocampus but not prefrontal cortex by environmental enrichment. **(a)** Expression of BDNF in the Hippocampus is reduced in RTT mice (light blue, n = 6 mice) compared to WT (red, n = 5 mice) and restored by environmental enrichment (dark blue, n = 4 mice). One-way ANOVA F (2,12) = 7.825 p= 0.006, Tukey’s post-hoc test, *p<0.05, **p<0.01. **(b)** Expression of BDNF in the Prefrontal Cortex is reduced in RTT mice (light blue, n = 6 mice), compared to WT mice (red, n = 5 mice) but could not be restored by environmental enrichment (dark blue, n = 4 mice). One-way ANOVA F (2,12) = 12.95 p = 0.001, Tukey’s post-hoc test, **p<0.01, ***p<0.001.

## DISCUSSION

In this study we describe morphological changes in prefrontal cortex connectivity in a mouse model of Rett Syndrome (**Figure 7**). Using genetically encoded fluorescent markers to track axonal projections from the prefrontal cortex, we carefully mapped their distribution across various cortical and subcortical regions. Our observations reveal several significant alterations in the projection patterns of prefrontal axons in heterozygous female *Mecp2^+/−^* mice. We demonstrate that the distribution of ACA axons in the cortex is altered in Rett model mice, with a stronger innervation of the somatosensory cortex and weaker targeting of the motor cortex (**Figure 3**). Consistent with the somatosensory cortex finding, the central network of prefrontal connectivity shows proportionately greater innervation (**Figure 4**). In addition, there is divergent distribution of prefrontal cortex axons between the ipsilateral and contralateral hemispheres, particularly in the most posterior regions of the cortex, so that the medial network of prefrontal connectivity shows proportionately reduced ipsilateral innervation in Rett model mice (**Figure 4**). Early exposure to environmental enrichment restores the deficits in motor coordination and elevated anxiety seen in *Mecp2^+/−^* mice (**Figure 1**) but does not affect the altered organization of axonal projections in the cortex. Moreover, the loss of MeCP2 lead to a reduction in BDNF expression, which is restored by environmental enrichment in the hippocampus but not in the prefrontal cortex (**Figure 6**). Our findings suggest that the effects of environmental enrichment can occur without large-scale changes in neuroanatomical projections (at least of prefrontal cortex neurons) and may occur either by changes in other focal structures (e.g., the hippocampus, cerebellum, or amygdala) or at finer (e.g., synaptic) levels than revealed by our study.

**Figure 7.**
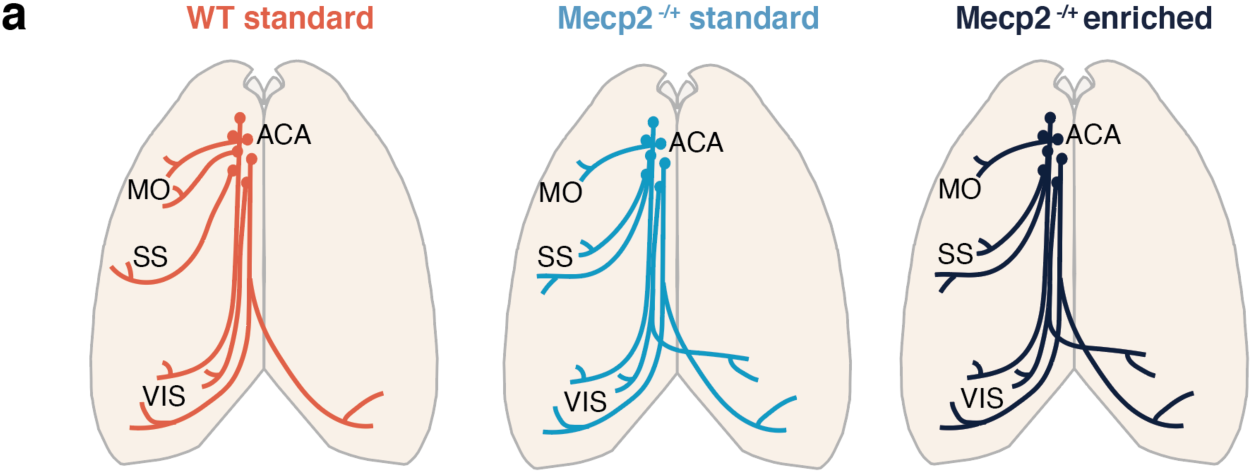
Alterations of ACA cortical projections in *Mecp2^+/−^* mice. **(a)** ACA axons in *Mecp2^+/−^* mice show increase in axon density in SS, loss in MO, and reduced ipsilateral dominance in posterior visual regions. This anatomical phenotype is not rescued by enriched housing. Complete list of abbreviations in **Supplementary Table 1.**

### Environmental enrichment rescues behavioral phenotypes of *Mecp2^+/−^* mice

*Mecp2^+/−^* female mice have progressive neurological symptoms, recapitulating disease progression seen in Rett patients, and are therefore ideal for pre-clinical studies on interventions to reduce or offset the progression (Samaco et al., 2013). Here we recapitulate previous work and show that behavioral phenotypes of *Mecp2^+/−^*female mice can be rescued with environmentally enriched housing in adolescence (Kondo et al., 2008, 2016; Lonetti et al., 2010). Fine motor skills, evaluated on the rotarod assay, of *Mecp2^+/−^* EH mice were equal to that of WT mice, while *Mecp2^+/−^* ST mice showed a severe deficit (**Figure 1c, d**). Similarly, the anxiety phenotype of *Mecp2^+/−^* mice was reversed with enriched housing, with *Mecp2^+/−^*EH mice spending more time in the center of the open field in comparison to both WT controls and *Mecp2^+/−^* ST mice (**Figure 1e**). Thus, environmental enrichment is a powerful means for rescuing important phenotypes of Rett Syndrome.

### Altered prefrontal innervation of somatosensory and motor cortex in *Mecp2^+/−^*mice

Previous work has shown that Rett model mice have neurophysiological and anatomical changes in the barrel cortex (Moroto et al., 2013; Lee et al., 2017; Morello et al., 2018) with weaker sensory evoked activity, reduced dendritic complexity and altered thalamocortical connectivity (Moroto et al., 2013; Lee et al., 2017). The primary motor cortex of Rett mice shows decreased spine density and disorganized axons with less myelination (Belichenko et al., 2009). In addition, the turnover of axonal boutons in the motor cortex, triggered by motor learning, is absent in a mouse model of *Mecp2* duplication syndrome (Ash et al., 2018). Thus, changes in MeCP2 expression have widespread effects on the functional and structural organization of somatosensory and motor cortex. Our finding of enhanced projections from ACA to somatosensory cortex but reduced projections to motor cortex in *Mecp2^+/−^* mice (**Figure 7**) is consistent with the view that these regions are particularly susceptible to MeCP2 dysregulation.

### Growth factors and axonal growth in *Mecp2^+/−^* mice

*Mecp2^+/−^* mice showed a reduction in BDNF expression, which was restored by environmental enrichment in the hippocampus but not in the prefrontal cortex (**Figure 6**). This is of particular interest as MeCP2 has been shown to influence the trafficking of BDNF levels in axons, which is essential for the development and maintenance of neuronal health (Altar et al., 1997; Roux et al., 2012). Local secretion of neurotrophins can influence the branching and stabilization of axons (Lewis et al., 2013). The stable axonal projection patterns across the two *Mecp2* groups could therefore be attributed to a consistent lack of neurotrophic factors, unaffected by enriched housing conditions. In addition, the developmental window of axonal growth is P1-P21 in mice (Lewis et al., 2013), indicating that the majority of axonal growth has already occurred once the *Mecp2^+/−^*mice are placed in enriched housing. Our enrichment timeline might therefore be implemented too late in development for large-scale axonal reorganization.

The expression level and secretion of BDNF is influenced by the activity of the local network (Brigadski and Leßmann, 2020). Environmental enrichment is known to increase the activity of the hippocampus and trigger neurogenesis in the region (Olson et al., 2006), consistent with upregulation of BDNF levels. In contrast, the decreased activity levels of frontal cortices observed in Rett model mice (Kron et al., 2012) might work against an environmentally-triggered rescue of BDNF levels in the prefrontal cortex.

In conclusion, our study is the first to identify specific, large-scale anatomical changes in prefrontal cortex projections in Rett model mice and confirms the role of environmental enrichment in restoring important phenotypes of Rett Syndrome. Our findings suggest that the mechanisms of such restoration do not involve reversing the large-scale anatomical changes but rather likely occur by engaging focal brain regions, and potentially involving functional changes at the level of synapses rather than widespread anatomical projections.

## AUTHOR CONTRIBUTIONS

All authors conceptualized the study. SÄR, GF and JH performed the surgeries. RL and JH performed the open field and rotarod assays. GF and JH performed the BDNF assays. JH ran the anatomical pipeline and behavioral tracking of open field data. SÄR analyzed and visualized the anatomical and open field data. RL analyzed and visualized the rotarod data. GF analyzed and visualized the BDNF data. MS provided funding and supervision. SÄR and MS wrote the manuscript with input from all authors.

## DATA AVAILABILITY

The data that support the findings of this study are available from the corresponding author upon reasonable request.

## FUNDING

This research was supported by Postdoctoral Fellowship WGF2020-0019 from the Wenner-Gren foundation and NIH grant 1K99EY035752 (SÄR), NIH grants R01MH085802, R01MH133066, R01NS130361, and The Picower Institute Innovation Fund (MS), the JPB Foundation Postdoctoral Fellowship (GF), NIH grant T32MH112510 (RL), and the MIT Brain and Cognitive Sciences Research Scholars Program (JH).

## CONFLICT OF INTEREST

The authors declare no competing interests.

## ACKNOWLEDGEMENTS

We thank Tatsuya Osaki for his assistance with the slide scanning microscope and image acquisition and Taylor Johns for administrative and technical assistance.

## Notes

### Competing Interest Statement

The authors have declared no competing interest.

