## Supplementary Material for "Persistent Disruptions in Prefrontal Connectivity Despite Behavioral Rescue by Environmental Enrichment in a Mouse Model of Rett Syndrome"

### Supplementary Figures

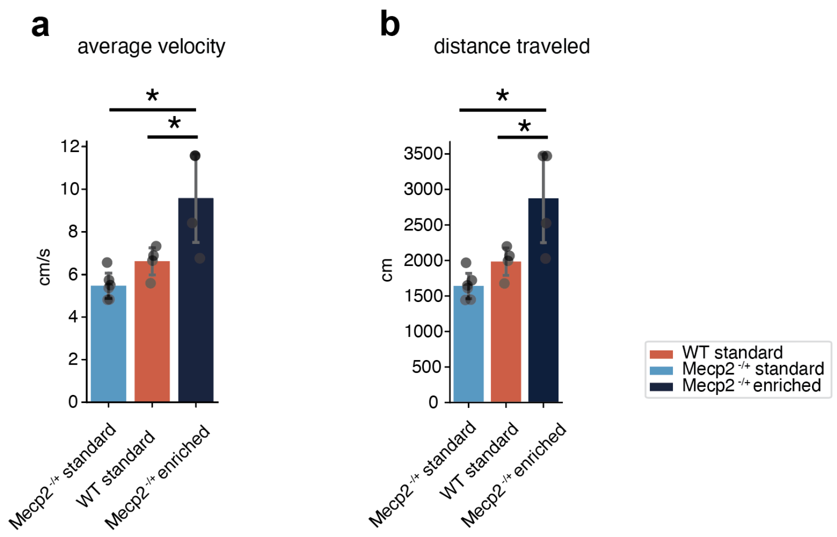


**Supplementary Figure 1. *Mecp2*$^{+/-}$ mice in enriched housing move more, and faster, than standard housed mice**

**(a)** Average velocity (cm/s) in the open field of WT standard housing (red, n=5 mice), Mecp2 standard housing (light blue, n=6 mice) and Mecp2 enriched housing (blue, n=4 mice) experimental groups. *p-value < 0.05, one-way ANOVA and Tukey's post-hoc adjustment. p-value = Mecp2 EH- WT: 0.028, Mecp2 EH- Mecp2 SH: 0.002.

**(b)** Cumulative distance traveled (cm) in the open field of WT standard housing (red, n=5 mice), Mecp2 standard housing (light blue, n=6 mice) and Mecp2 enriched housing (blue, n=4 mice) experimental groups. *p-value < 0.05, one-way ANOVA and Tukey's post-hoc adjustment. p-value = Mecp2 EH- WT: 0.028, Mecp2 EH- Mecp2 SH: 0.002. Bars represent mean values; error bars are standard error and points represent individual animals. Complete list of abbreviations in **Supplementary Table 1.**

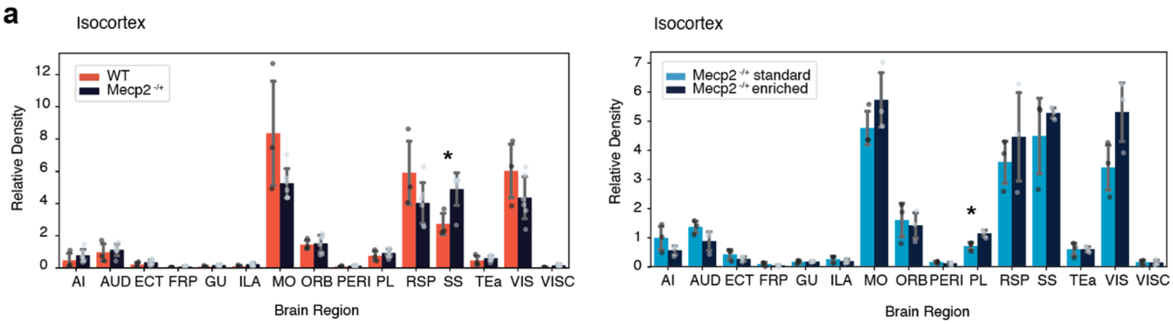


**Supplementary Figure 2. Relative Density of ACA axons per cortical region in WT and *Mecp2*$^{+/-}$ mice**

**(a)** (Left) Relative Density of ACA axons per cortical region out of all Isocortex in WT (red, n=3 mice) and Mecp2 (dark blue, n=6 mice) mice. *p-value < 0.05, unpaired Student's t-test. SS p-value: 0.020. (Right) Relative Density of ACA axons per cortical region out of all Isocortex comparing Mecp2 standard housing (light blue, n=3 mice) and Mecp2 enriched housing (blue, n=3 mice) experimental groups. *p-value < 0.05, unpaired Student's t-test. SS p-value: 0.023. Related to Figure 1. Complete list of abbreviations in **Supplementary Table 1.**

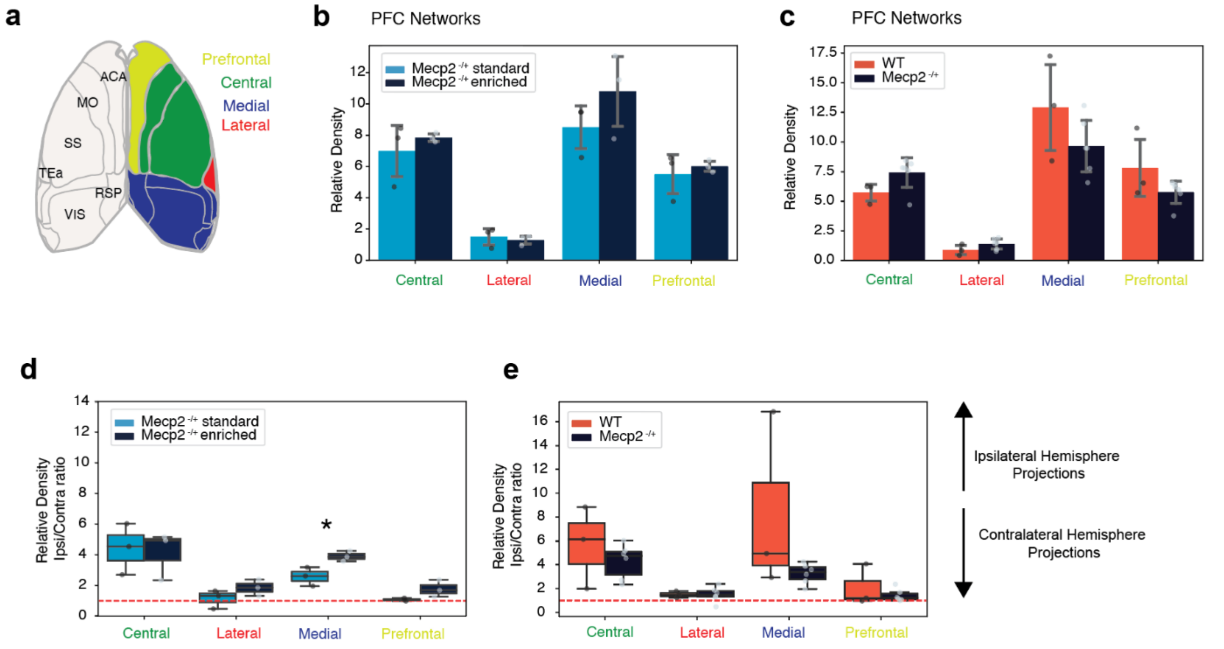


**Supplementary Figure 3. Relative Density of ACA axons per PFC network in WT and *Mecp2*$^{+/-}$ mice**

**(a)** Schematic of cortical regions participating in each PFC network. Prefrontal: FRP, PL, ILA, ORB, ACA, MOs, AId, AIv. Lateral: AIp, GU, VISC, TEa, PERI, ECT, ENT. Central: MOp, SS. Medial: PTLp, RSP, VIS, AUD.

**(b)** Relative Density of ACA axons for brain regions participating in each PFC network for Mecp2 standard housing (light blue, n=3 mice) and Mecp2 enriched housing (blue, n=3 mice) experimental groups. Bars represent mean values; error bars are standard deviation and points represent individual animals.

**(c)** Relative Density of ACA axons for brain regions participating in each PFC network in WT (red, n=3 mice) and Mecp2 (dark blue, n=6 mice) mice.

**(d)** Ratio of Relative Density between ipsilateral and contralateral hemispheres for ACA axons innervating each PFC network for Mecp2 standard housing (light blue, n=3 mice) and Mecp2 enriched housing (blue, n=3 mice) experimental groups. *p-value < 0.05, unpaired Student's t-test. Medial p-value: 0.031.

**(e)** Ratio of Relative Density between ipsilateral and contralateral hemispheres for ACA axons innervating each PFC network in WT (red, n=3 mice) and Mecp2 (dark blue, n=6 mice) mice. Each boxplot represents the quartiles of values, while the whiskers extend to show the rest of the values within 1.5 times the interquartile range. Points represent individual animals. Complete list of abbreviations in **Supplementary Table 1.**

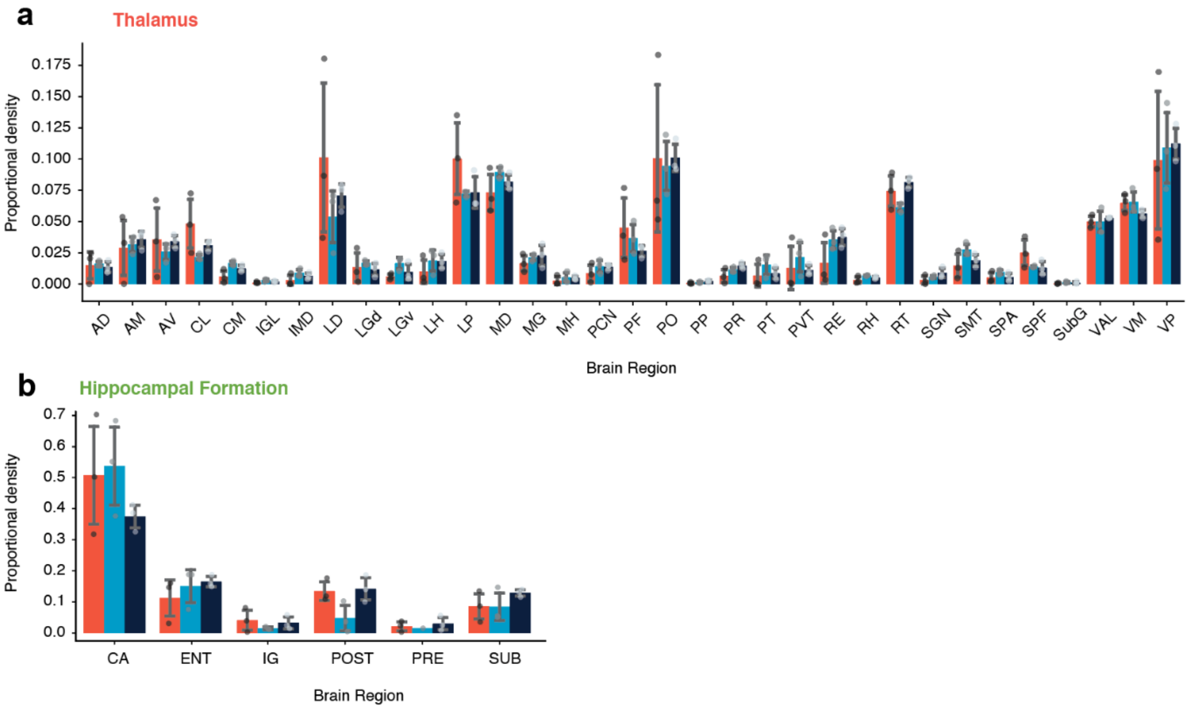


**Supplementary Figure 4. Subcortical projections of ACA neurons**

**(a)** Proportional density of ACA axons in Thalamic nuclei for WT standard housing (red, n=3 mice), Mecp2 standard housing (light blue, n=3 mice) and Mecp2 enriched housing (blue, n=3 mice) experimental groups.

**(b)** Proportional density of ACA axons in the Hippocampal Formation for WT standard housing (red, n=3 mice), Mecp2 standard housing (light blue, n=3 mice) and Mecp2 enriched housing (blue, n=3 mice) experimental groups. Bars represent mean values; error bars are standard deviation and points represent individual animals. Complete list of abbreviations in **Supplementary Table 1.**

**Supplementary Table 1. Nomenclature**

**ISO Isocortex**

MO Motor cortex

MOp Primary motor area

MOs Secondary motor area

SS Somatosensory areas

SSp Primary somatosensory area

SSs Secondary somatosensory area

ILA Infralimbic area

GU Gustatory areas

VISC Visceral area

AUD Auditory areas

VIS Visual areas

ACA Anterior cingulate area

PL Prelimbic area

ORB Orbital area

AI Agranular insular area

RSP Retrosplenial area

TEa Temporal association areas

PERI Perirhinal area

FRP Frontal Pole

**HPF Hippocampal formation**

CA Ammon's Horn

IG Induseum griseum

ENT Entorhinal area

POST Postsubiculum

PRE Presubiculum

SUB Subiculum

**CTXsp Cortical Subplate**

**CNU Cerebral nuclei**

CP Caudoputamen

ACB Nucleus accumbens

LS Lateral septal nucleus

GPe Globus Pallidus, External segment

GPi Globus Pallidus, Internal segment

SI Substantia innominata

MA Magnocellular nucleus

NDB Diagonal band nucleus

**TH Thalamus**

VAL Ventral anterior-lateral complex of the thalamus

VM Ventral medial nucleus of the thalamus

VP Ventral posterior complex of the thalamus

SPF Subparafascicular nucleus

SPA Subparafascicular area

LP Lateral posterior nucleus of the thalamus

PO Posterior complex of the thalamus

AV Anteroventral nucleus of the thalamus

LD Lateral dorsal nucleus of the thalamus

MD Mediodorsal nucleus of the thalamus

SMT Submedial nucleus of the thalamus

RE Nucleus of reunions

CL Central lateral nucleus of the thalamus

PF Parafascicular nucleus

RT Reticular nucleus of the thalamus

PP Peripeduncular nucleus

POL Posterior limiting nucleus of the thalamus

SGN Suprageniculate nucleus

AM Anteromedial nucleus of the thalamus

AD Anterodorsal nucleus of the thalamus

IMD Intermediodorsal nucleus of the thalamus

PR Perireuniens nucleus

PVT Paraventricular thalamic nucleus

PT Paratenial nucleus

RH Rhomboid nucleus

CM Central medial nucleus of the thalamus

PCN Paracentral nucleus of the thalamus

**HY Hypothalamus**

**MB Midbrain**

SNr Substantia nigra, reticular part

VTA Ventral tegmental area

MRN Midbrain reticular nucleus

PAG Periaqueductal gray

SC Superior Colliculus

APN Anterior pretectal nucleus

**HB Hindbrain**
